# MYB68 regulates radial endodermal differentiation and suberin patterning

**DOI:** 10.1101/2024.05.09.593291

**Authors:** Leonie Kraska, Josep Mercadal Melia, Ryohei Thomas Nakano, David Molina, Pau Formosa-Jordan, Laura Ragni, Tonni Grube Andersen

## Abstract

Roots are composed of concentric tissue layers that embrace the centrally localized vasculature. Of these layers, particularly the endodermis stands out as it contains barriers that facilitate selective uptake across the plasma membrane. In mature root regions, endodermal cells undergo additional differentiation and become coated with suberin, a hydrophobic polymer that blocks membrane transport and seals off the inner root parts. Intriguingly, individual cells adjacent to the water-conducting xylem remain unsuberized. These are termed “passage cells”, based on the assumption that they facilitate radial vascular access in a zone which is otherwise impenetrable. The identity of passage cells remain unknown, but their existence suggests that distinct identities and developmental trajectories exist within the second differentiation of the endodermis. In this study, we investigate this in the model plant *Arabidopsis thaliana*. Our work identifies a genetic regulator that controls pole-specific endodermal differentiation and tissue-forming divisions connected to passage cells. Through a number of analyses, we provide spatiotemporal insights into suberization, establish a framework for radial organization of the endodermis and highlight putative function(s) of passage cells. Combined our findings illustrate how multi-dimensional developmental proceses integrate with environmental inputs in order to provide distinct cellular functions within root tissues.

## Introduction

The organization of root cell layers is crucial for selective filtering of solutes translocated between the above- and below-ground plant parts. In *Arabidopsis thaliana* (hereafter Arabidopsis), the outer tissues of the primary root comprise the soil-facing epidermis and two underlying layers: the cortex and the endodermis, which combined constitute the ground tissues (Dolan, et al., 1993). The endodermis has historically received the most attention, as it contains apoplastic diffusion barriers that are key for uptake of minerals and nutrients into the xylem (Geldner, 2013). In contrast to the radially symmetric outer layers, the central vasculature is organized in opposing xylem and phloem poles faciliating upward water/mineral and downward sugar transport, respectively. Anatomically, the endodermis overlays the vasculature and the individual cells within its circumference largely align with these vascular poles. Thus, endodermal cells can be divided into those that are xylem pole-associated (XPE), phloem pole-associated (PPE) or situated inbetween (non-pole-associated, NPE). No distinct identity or function has yet been ascribed to these cells, but after about one week of growth, sporadic formative divisions in the early XPE cells initiate an additional ground tissue layer called the middle cortex (MC) (Baum, et al., 2002). The function and genetic program underlying MC cells are not fully understood, but their existence suggests that XPE cells contain specialized developmental information related to their xylem-associated position (Baum et al. 2002; (Paquette and Benfey, 2005). Although distinct from MC, a second cortex cell layer has also been described to occur close to the hypocotyl in young roots (Scheres, et al., 1994).

As the root grows and cells exit the meristematic root tip, the endodermis undergoes two stages of differentiation before dying off during periderm formation (Wunderling, et al., 2018; Geldner, 2013). The first involves deposition of the lignin-based apoplastic barrier known as the Casparian strip (CS) (Caspary, 1865). The CS prevents extracellular diffusion and forces movement of solutes across the plasma membrane (Priestley and North, 1922). The second stage is characterized by deposition of the hydrophobic polymer suberin in the form of a lamellae-like structure across the entire cell surface (Franke, et al., 2005; Sitte, 1959). Suberin blocks transport across the plasma membrane and thereby seals off the vasculature from its external surroundings (Andersen, et al., 2015).

In 5-6-day-old Arabidopsis seedlings, the differentiation towards suberization occurs approximately 15-20 cells after formation of the Casparian strip (CS) and follows a stereotypical pattern along the longitudinal root axis (Alassimone, et al., 2010). The decision to suberize occurs sporadically and slightly earlier in the PPE than XPE and NPE (Andersen, et al., 2018). This gives ground to a so-called “patchy zone” of about 10-15 endodermal cells along the longitudinal axis of the root where not all cells across the endodermal circumference are suberized. After this transition, a “fully suberized” zone is established, covering the remaining endodermis to the root-hypocotyl junction. Intriguingly, and in line with distinct radial functions within the endodermis, a few individual cells in the XPE and NPE remain unsuberized (Holbein, et al., 2021; Andersen, et al., 2018; Peterson and Enstone, 1996; Kroemer). These are referred to as endodermal “passage cells” (PCs), as their lack of suberization is assumed to provide a low-resistance radial flow path for nutrients and water into the xylem in an otherwise isolated part of the root (Peterson and Enstone, 1996; Kroemer). Although PCs were identified over a century ago, and have been observed in many plant lineages (Holbein, et al., 2021), genetic insights into their function and development have only recently begun to emerge. In Arabidopsis, the first steps toward PC formation are initiated in the meristematic XPE by radially organized hormonal signal mechanisms emanating from the developing xylem (Andersen, et al., 2018; Mähönen, et al., 2006). However, the genetic network downstream of this decision as well as mechanisms that initiate the differentiation towards suberization remain unknown.

In this study, we set out to find factors that control endodermal differentiation towards suberization with focus on elucidating radial specification. Our work identifies MYB68, a homolog of the CS master regulator MYB36, as an important player for suberization of the XPE and NPE endodermal cell files. Through transcriptional profiling we highlight that MYB68 is involved in a genetic network that regulate XP-associated ground tissue differentiation. We find that MYB68 likely functions both in mature and young endodermal cells, where it is implicated in an age-dependent mechanism that determine formation of MC and PCs in the XPE. Combined, this brings about new insights into root cell identity establishment and provides tools for deeper understanding of spatiotemporal patterning and tissue functions.

## Results

### MYB68 influences endodermal suberization

To start our investigation, we focused on a subclade of MYB TFs that includes the master regulator of CS formation MYB36 (Kamiya, et al., 2015; Liberman, et al., 2015). We hypothesized that within this clade, other members may play roles associated with endodermal differentiation. This subfamily comprises six members: MYB36, MYB37, MYB38, MYB68, MYB84, and MYB87, for which we recently established homozygous knockout (KO) mutant lines (Molina, et al., 2024). For MYB68, a previous study identified a KO allele in the Landsberg ecotype (Feng, et al., 2004) and we therefore named our alleles *myb68-2* and *myb68-3*. To assess whether mutants of this clade of MYBs are affected in endodermal development, we examined functionality of the CS and measured suberin patterning in 6-day-old roots. Consistent with previous findings (Kamiya, et al., 2015; Liberman, et al., 2015), specifically *myb36-2* exhibited a dysfunctional apoplastic barrier, evident by a significant increase in propidium iodide (PI) penetration into the vasculature (Naseer, et al., 2012). m*yb36-2* also displayed an early suberization onset, which is associated with its impaired CS function (Doblas, et al., 2017; Kamiya, et al., 2015; Liberman, et al., 2015) (**Figure 1A and S1A**). Interestingly, besides *myb36-2*, only *myb68* alleles exhibited changes in relation to suberization (**Figure 1B and S1B**). This was manifested as a significant increase in the relative size of the patchy suberized zone, which appeared to be due to absence of suberin in neighbouring cells (**Figure 1D**). To investigate whether this was related to a general defect in the ability to synthesize or deposit suberin, we exposed *myb68* roots to 1 μM ABA for 2 days to induce endodermal suberin deposition (Barberon, et al., 2016). This resulted in a significantly earlier onset of coherent suberin deposition in both Col-0 and *myb68* roots (**Figure 1C and S1B**). Combined, these findings therefore identifiy MYB68 to function in a genetic network that controls suberin patterning in the endodermis rather than its direct biosynthesis or deposition.

**Figure 1.**
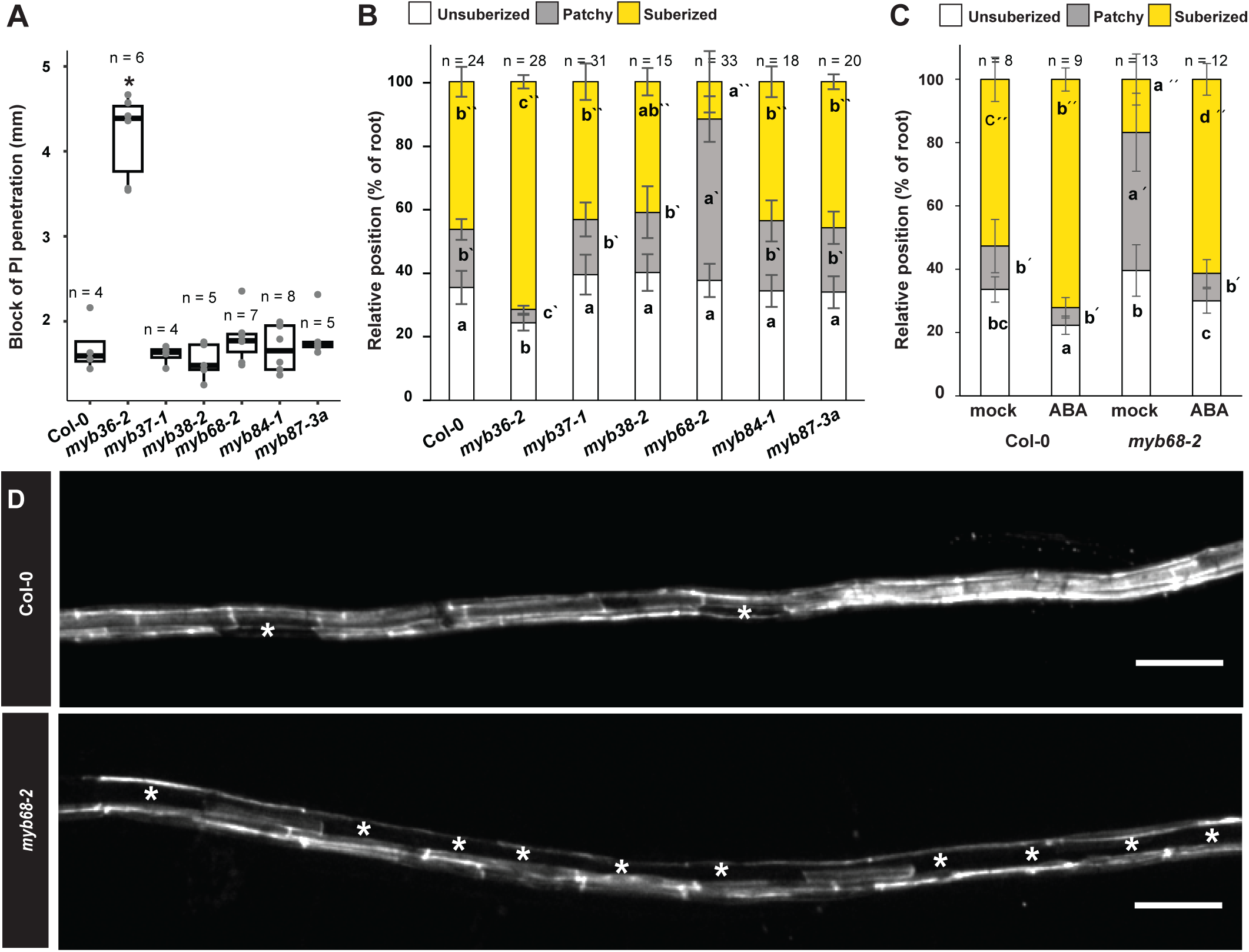
| Formation of endodermal barriers in MYB36-clade mutants. (A) Functional analysis of CS by measuring onset of propidium iodide (PI) diffusion blockage into the stele (Naseer, et al., 2012). Asterisks depict significant differences to WT according to the unpaired Wilcoxon test. B) and C) Suberization patterning in 6-day-old roots stained with Fluorol Yellow 088 (FY). D) Confocal maximum projection of the suberizing zone (aprox 60% relative position) of Col-0 and myb68-2 root stained with FY, asterisks mark individual unsuberized cells. Individual letters depict significance (P<0.05) according to a Kruskal Wallis test with a post hoc Nemenyi test. Letters without a prime refer to unsuberized zones, letters with one prime refer to patchy zones and letters with two primes refer to fully suberized zones. Scale bar represents 100 μm. ABA: Abscic acid, PI: Propidium Iodide.

### Suberin deposition displays cell file-specific MYB68-dependent dynamics

To investigate if the changed pattern of suberization is related to the vascular-associated positions of the endodermis, we measured the suberin status of the individual radial cells (XPE,NPE and PPE) in 6-day-old roots (**Figure 2A**). By plotting the cumulative sum of suberized cells against their relative position along the longditudinal axis, we could assess the suberization continuity of the individual cell files from the onset until the root-hypocotyl junction (for detailed description see materials and methods). In both Col-0 and *myb68-2* roots, PPE cell lineages gave rise to a linear behavior, although deviations could be observed in *myb68-*2 due to the presence of a few unsuberized cells (**Figure 2A**). For NPE cell files, both genotypes displayed a bilinear trend, reflecting the patchy and fully suberized zones respectively. In XPE, and to a lesser extend NPE files, the suberization frequency was reduced in *myb68-2* roots when compared to Col-0 (**Figure 2A**). MYB68 therfore appears to primarily influence suberization in XPE and NPE cell files, and thereby play a role in regulation of the patchy zone. One intriguing observation was that Col-0 roots consistently had a decrease in the number of suberized cells in XPE and NPE files at the upper end of the roots close to the hypocotyl (**Figure 2A**), suggesting the existence of an additional unsuberized area at this developmental stage.

**Figure 2.**
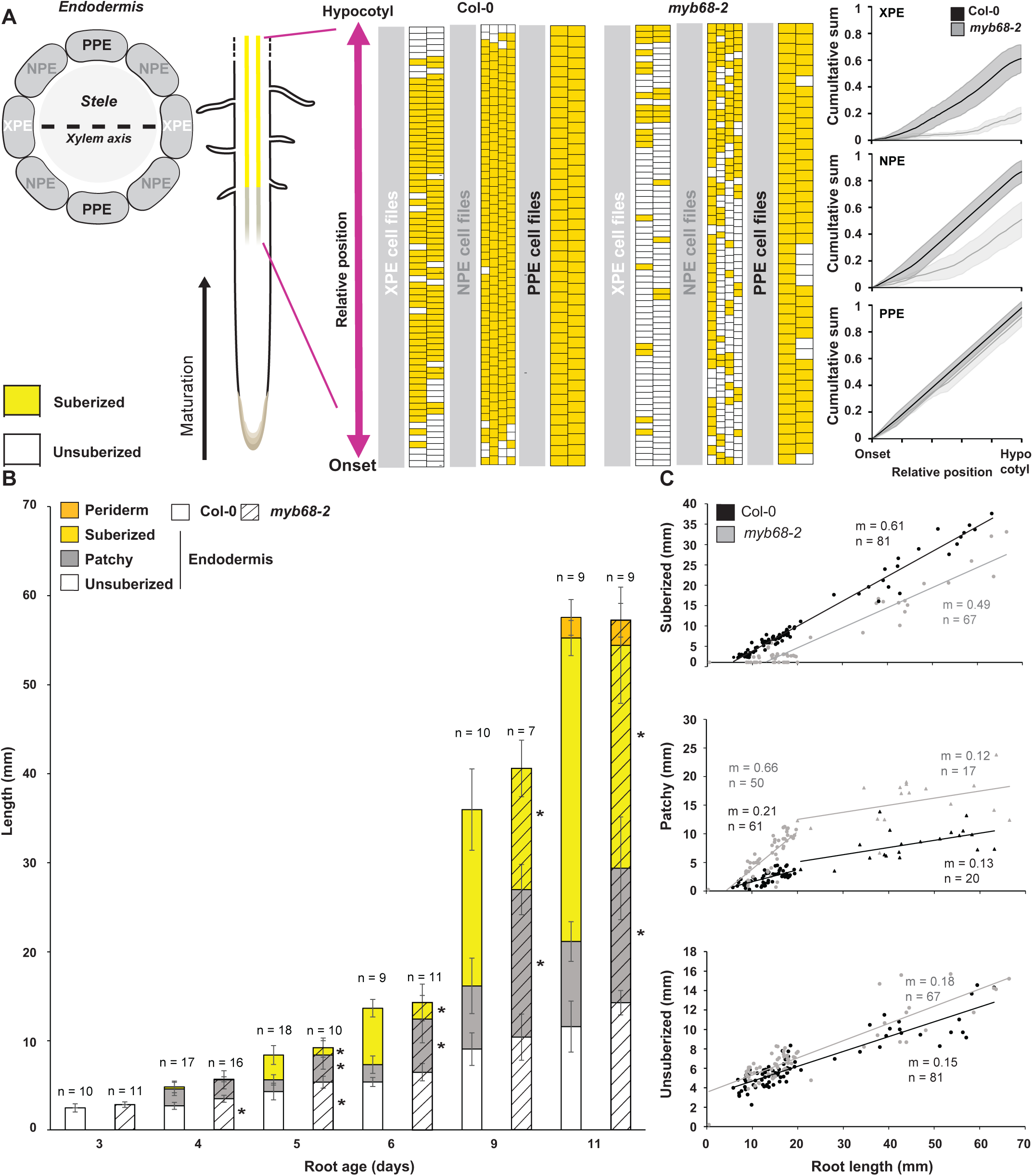
| Spatiotemporal analysis of endodermal suberization patterns. A) Analysis of radial suberin patterning of roots stained with Fluorol Yellow 088 (FY). The cumulative sum of each cell file along the upper 50% of the root was plotted for WT and myb68-2against the position on the longitudinal axis (materials and methods). The lines in graphs depict the average of cell files across 3 individual roots and shading the standard deviation. B) Time-course analysis of suberin patterning in roots from 3 to 11 days of age stained with FY. Asterisks indicate significant differences between Col-0 and myb68-2 at each time point, and in each zone via an unpaired Wilcoxon text (* p value < 0.05). C) Measurement of the relationship between suberin zones and root length of individual roots from 3- to 11-day-old roots. Lines depict a linear regression fit with calculated values for the slopes (m) (see Table S1). XPE: Xylem pole associated endodermis; PPE: Phloem pole associated endodermis; NPE: non-pole associated endodermis.

We next measured the progression of suberization from 3- to 11-days-old roots, which covers the entire endodermal life span (Serra, et al., 2022; Wunderling, et al., 2018). In both Col-0 and *myb68-2*, the fully suberized area correlated positively with root length (**Figure 2B and 2C**). This was expected, as both the aging endodermal cells and the periderm are suberized and comprise an increasing proportion as the root grows (Wunderling, et al., 2018). However, the length of unsuberized and patchy areas also increased with age (**Figure 2B and 2C**), with the latter stabilizing when roots reached a length of approximately 20 mm (**Figure 2C**). This was suprising as these zones are assumed to be coordinated with root growth and therefore display a stable proportion of the root over time. Thus, these findings suggest that age-dependent mechanism(s) may underlie the decision suberize. In line with a role of MYB68 in the networks regulating this, *myb68-2* roots displayed a delay in full suberization, but with a similar patterning rate and onset as Col-0 (**Figure 2B, 2C and Table S1**).

### MYB68 controls occurrence of endodermal passage cells

As the observed decrease in suberized cells in XPE cell files of *myb68-2* roots could reflect a change in PC formation, we investigated if MYB68 influences the establishment of PCs. These cells are typically defined by their lack of suberin and therefore indistinguishable from unsuberized cells, rendering them impossible to access in this connecton. However, several genes have been described to be associated with PC, including the *phosphate exporter homologue* 3 (*PHO1;H3*) (Andersen, et al., 2018). We therefore deployed a transcriptional marker (*pPHO1;H3: NLS 3x mVenus*) for this gene and measured how often we could observe expression in the endodermis which is normally undergoing suberization. Indeed, consistent with PC-association, mainly cells in the XPE and NPE of the suberized zone of wildtype roots displayed *PHO1;H3* activity. Intriguingly, and in line with the reduced suberization, this pattern was exaggerated in *myb68-2* roots (**Figure 3A and 3B**). Thus, MYB68 may be involved in repression of PC occurence. In support of this, germination of *myb68-2* on PC-suppressive concentrations of the artificial cytokinin benzylaminopurine (BAP) (Andersen, et al., 2018), led to a dose-dependent increase of suberization as well as a corresponding decrease in PC occurrence (**Figure 3C and 3D**).

**Figure 3.**
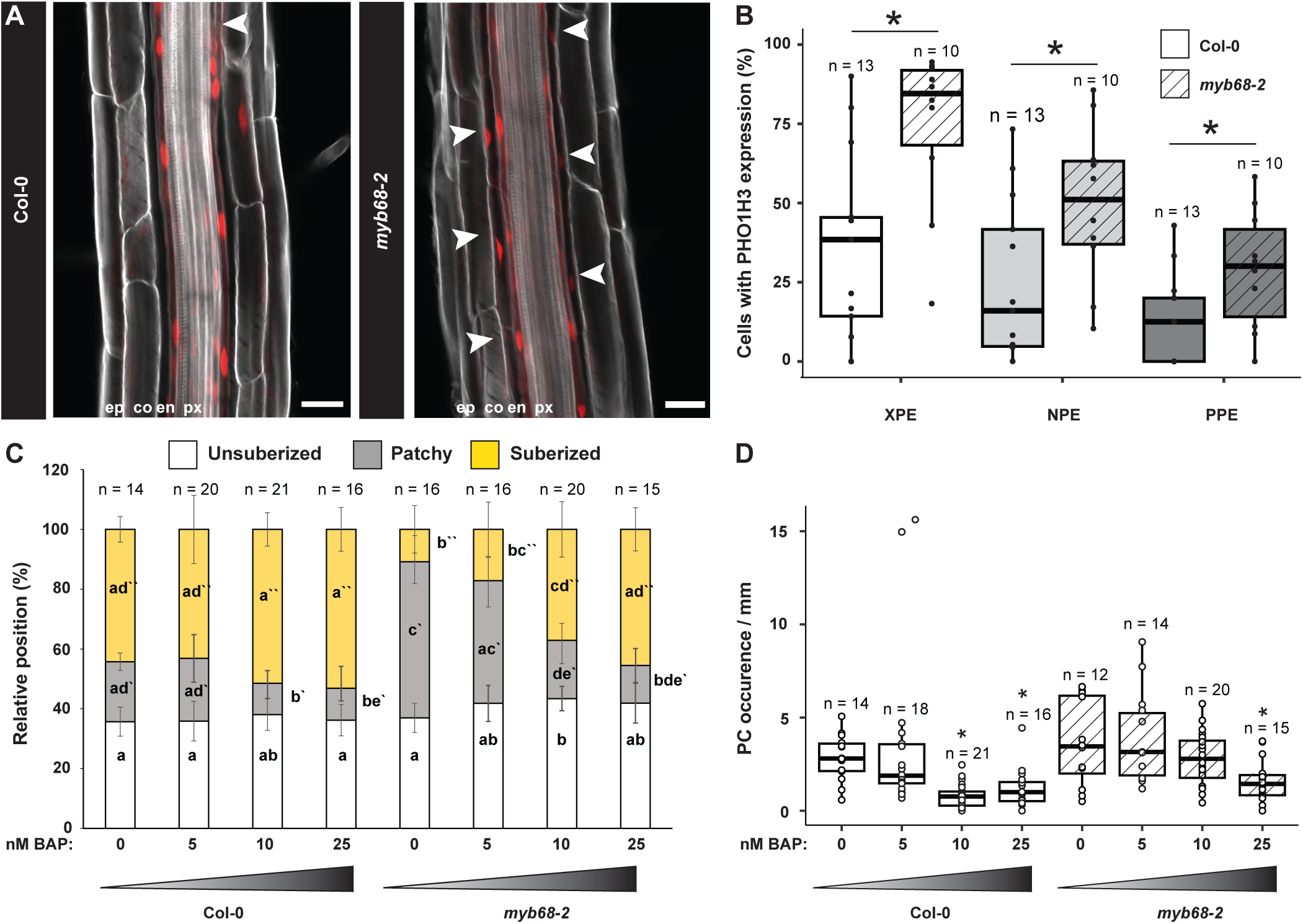
| Investigation of passage cell occurrence in the *myb68-2* mutant. A) Representative root of a transcriptional marker line reporting activity of the promoter region of a PHO1;H3 (pPHO1;H3: NLS 3xmVenus) (Andersen, et al., 2018). Image originates from the zone of continuous suberization in a 6-day-old root. Arrowheads highlight endodermal cells with signal. PHO1;H3 activity is depicted in red and the cell walls were stained with Calcoflour white (grey) according to (Ursache, et al., 2018). B) Proportion of endodermal cells with expression of PHO1;H3 in the xylem pole-associated endodermis (XPE), non-pole associated endodermis or phloem pole-associated endodermis (PPE) in the upper 50% of the root. Each data point represents the percentage within each analyzed root. Stars indicate significant difference between PHO1;H3 activity in WT (pGPAT5:mCitrine-SYP122) (Andersen, et al., 2018) and myb68-2 background lines in each longitudinal position according to unpaired Wilcoxon test (* p value < 0.05). C) Suberization pattern in roots grown in the presence of increasing amounts of the artificial cytokinin 6 Benzyl-aminopurine (BAP). Individ-ual letters depict significance (P<0.05) according to a Kruskal Wallis test with a post hoc Nemenyi test. Primes refer to zones of suberiza-tion status. D) Passage cell (PC) occurrence in roots from C. Asterisks indicate significant differences between mock- and BAP-treated samples in an unpaired Wilcoxon text (* p value < 0.05). XPE; Xylem pole associated endodermis, PPE; Phloem pole associated endodermi**s, NPE; Non-pole associated endodermis**

### A subset of transporter-encoding genes associate with passage cells

Based on the apparent increase in PC occurrence of the *myb68-2* mutants, we next set out to probe if this could give insights into their transcriptional identity. We reasoned that genes associated with PCs would have to fulfill two criteria: 1) have upregulated expression in *myb68-2* when compared to Col-0 and 2) display Col-0-like expression levels upon BAP treatment, as this restored *myb68-2* suberization back to a similar pattern as observed in mock-treated Col-0 (**Figure 3C**), assuming that occurrence of PC was similarly restored. 296 genes were upregulated in *myb68-2*, of which 66 were repressed upon BAP treatment (**Figure 4A, Table S2 and S3**). This subset did not include *PHO1;H3,* which may be due to their PC-associated expression being masked by strong expression in the stele (Hamburger 2002, Andersen 2018). Thus, our selection criteria probably define more specific PC-associated components and possibly overlook genes with wider expression patterns. Despite this limitation, a gene ontology (GO) term analysis of the 66 PC-associated candidates (Zhou, et al., 2019) showed a significant enrichment in functions such as “inorganic cation transmembrane transport” “response to gibberellin” and “regulation of cell communication and signaling” (**Figure 4B**). This is in line with previously proposed roles of PCs in nutrient homeostasis as well as biotic communication (Holbein, et al., 2021) and, therefore likely contain new candidates for PC-associated expression.

**Figure 4.**
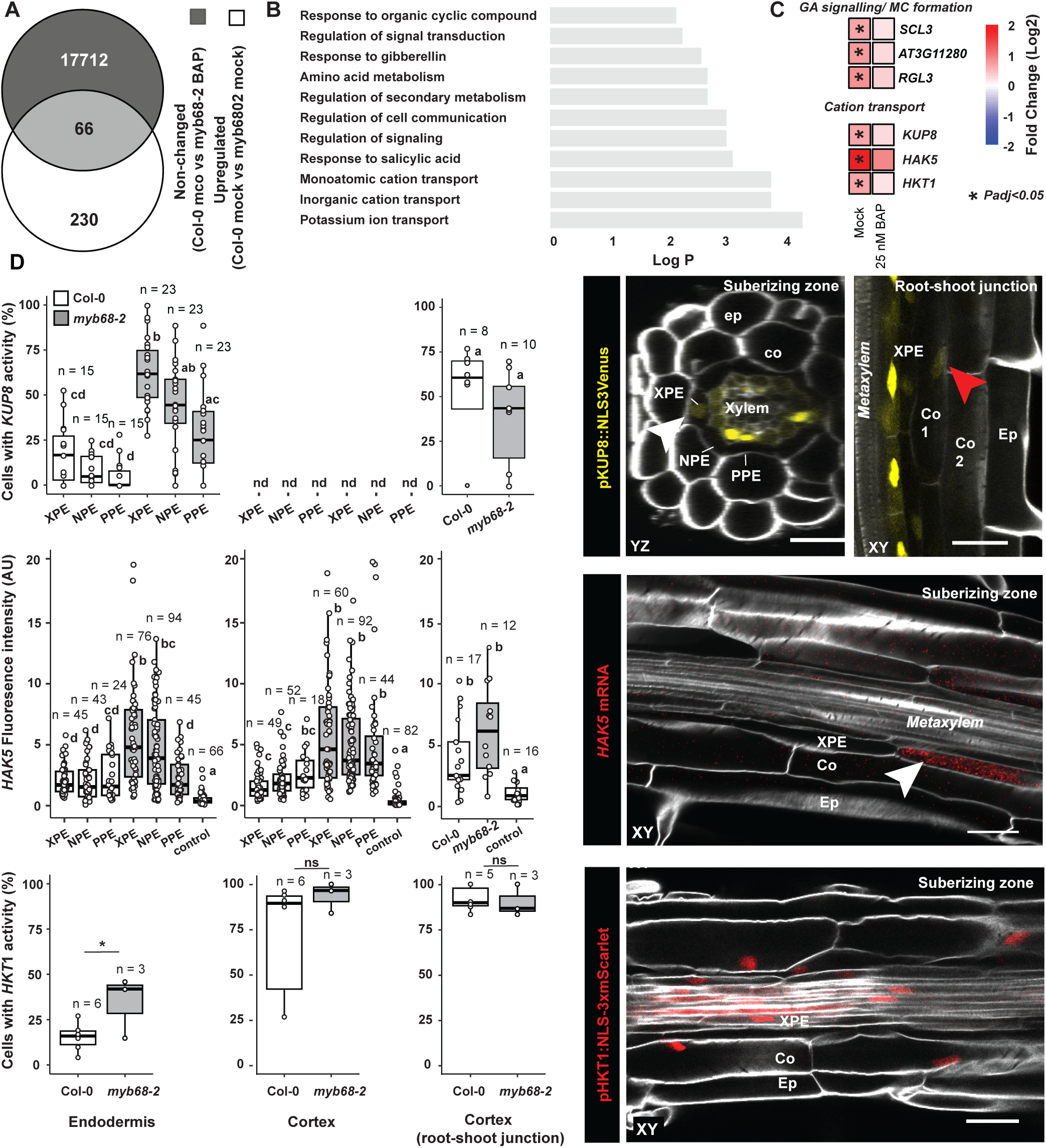
| Passage cell-associated transcriptional analysis. A) Overlap of upregulated genes in 6-day-old Col-0 vs myb68-2 roots (padj <0.05) and genes with non-significant changes in Col-0 vs. myb68-2 when germinated on BAP (25 nM 6-Benzyl-aminopurine). B) GO-term analysis of the 66 overlapping genes from A. C) Heatmap of selected gibberellic acid (GA) and middle cortex (MC) - associated genes as well as cation transporters in the comparisons from A (all genes in Table S4 and S5). D) Transcriptional activity of gene expression in the suberizing zone as well as in the inner cortex close to the root-shoot junction. Expression analysis of KUP8 and HKT1 in Col-0 and myb68-2 was performed using fluorescent transcriptional marker lines (expressing pKUP8:NLS3xVenus or pHKT1:NLS-3xmScarlet respectively). Each data point represents percentage of cells with expression in individual roots minimum 7 cells per root). For HAK5 mRNA, fluorescence in situ hybridization was employed. As negative control either a probe specific for a bacterial gene (dapB, Bacillus subtilis) with the corresponding amplifier or only the amplifier used for HAK5 without a probe were used. (see materials and methods). Individual letters show significance according to a Kruskal Wallis test with a post hoc Nemenyi test. XPE; Xylem pole associated endodermal cells, PPE; Phloem pole associated endodermal cells, NPE; non-pole-associated endodermal cells. Ep; Epidermis, Co; Cortex, En; Endodermis, nd; not detected, ns; not significant. Scale bars represent 25 µm.

Within the transport-related GO-term, we identified the two potassium (K) transporters *POTASSIUM UPTAKE 8* (*KUP8) and HIGH AFFINITY K+ TRANSPORTER 5 (HAK5)* (Osakabe, et al., 2013; Gierth, et al., 2005) as well as the Arabidopsis sodium (Na) transporter *HIGH-AFFINITY K+ TRANSPORTER* 1 (*HKT1)(Mäser, et al., 2002)* (**Figure 4C, Table S4**). To probe if the expression of these are indeed associated PCs, we created promoter-based transcriptional marker lines or performed whole-mount hybridization chain reaction fluorescent *in situ* hybridization (HCR-FISH) (Oliva, et al., 2022). Interestingly, in the suberizing part of 6-day-old Col-0 roots, *KUP8* showed activity in both endodermis, pericycle and specifically the inner cortex cells close to the hypocotyl where two layers had formed **(Figure 4D and S2A)**. *HAK5* mRNA was localized in endodermis and most cortex cells (including those close to the hypocotyl) (**Figure 4D, S2C)**, whereas *HKT1* displayed activity across all the investigated tissues (**Figure 4D, S2B and S2D**). Upon quantification, the expression of *KUP8* showed a bias towards XPE ans NPE cells and was significantly increased in the *myb68-2* mutant across all endodermal cell files (**Figure 4D**). For *HAK5*, although this expression was equally present across cell files in Col-0, specifically XPE, NPE and similar cortex cell files displayed a significant increase of expression in *myb68-2*. For *HKT1*, only endodermal expression was significantly increased in *myb68-2* **(Figure 4D, S1C and S3D)**. Combined, these analyses provids evidence for candidate genes that are likely associated with PC function and highlight a putative role of these elucive cells in cation homeostasis.

### MYB68 represses endodermal division in the apical meristem

Within the subset of genes with PC-correlated expressional behavior, we additionally observed several related to gibberelin signaling as well as the MC-connected *SCARECROW LIKE 3* (SCL3) *SCL3 (Yoshida, et al., 2014; Zhang, et al., 2011)* (**Figure 4C**). Thus, MYB68 may additionally influence hormonal responses associated with the formation of MC cells in the meristematic XPE file. To analyse this, we grew plants for 6 days on agar plates with a mesh filter, as this gave a more consistent establishment of MC under our growth conditions. In line with our transcriptomic analysis, *myb68-2* and *myb68-3* mutants displayed a significant increase in MC occurence (**Figure 5A, 5B and S1D**), suggesting that these cell types in XPE files may be repressed by MYB68.

**Figure 5.**
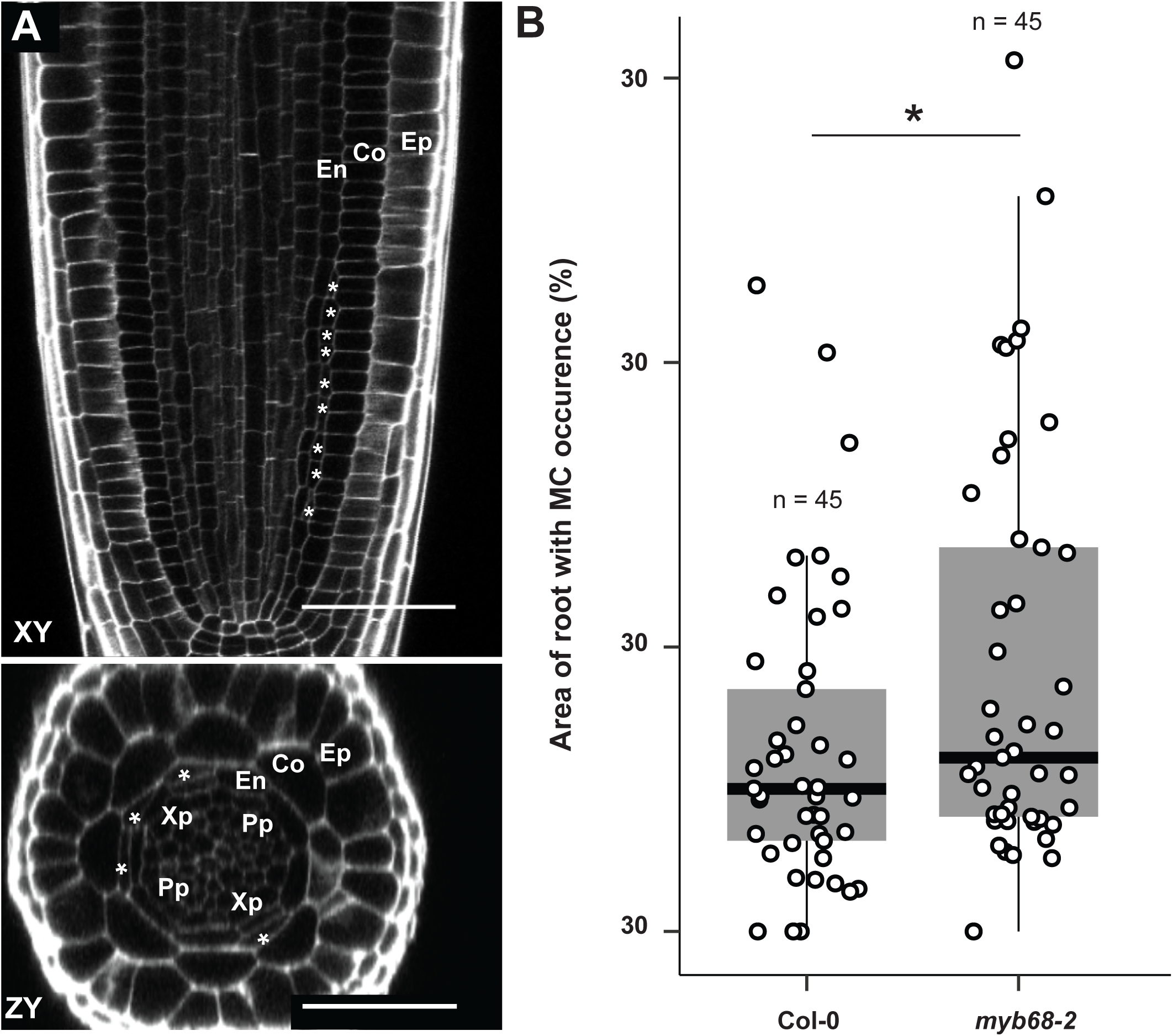
| Middle cortex cell occurence in 6-day-old *myb68-2* roots. A) Representative image of a longitudinal (XY) and transversal (ZY) section of a 6-day-old myb68-2 meristem with middle cortex (MC). Cellwalls were stained with Calcoflour white (grey) according to (Ursache, et al., 2018). Asterisks indicate MC cells. B) Measurement of MC occurrence in 6-day-old roots. The graph depicts the percentage of root with MC occurrence measured by defining the distance from the first-occurring MC to the root tip. Ep; Epidermis, Co; Cortex, En; Endodermis, Pp; Phloem pole, Xp; Xylem pole. I B) Asterisks indicate significant difference to Col-0 according to unpaired Wilcoxon text (* p < 0.05). Scale bars represent 50 µm.

### *MYB68* is expressed in both differentiated and meristematic tissues

The influence of MYB68 on MC cells is intriguing, as this proposes a role for MYB68 in integrating suberization patterning and periclinal division of the XPE cells. In order to determine if this is a direct function, we set out to characterize the expression pattern of MYB68. For this, we constructed fluorescence-based reporter lines suitable for investigating both transcriptional and translational expression. Our markers used the entire intergenic region upstream of *MYB68* to drive the expression of a nuclear-localized fusion reporter (NLS 3xmVenus) or a complementing genomic DNA fragment encoding MYB68 fused N-terminally to a GFP reporter (**Figure S3D**). In support of a local effect on suberization, both reporters displayed activity in the endodermis and in other vascular-associated tissues in the root parts corresponding to the suberization zone (**Figure 6A and 6F**). Our translational reporter showed accumulation of GFP-MYB68 in the endodermis and vascular tissues in the elongation zone (**Figure 6F**). Intriguingly, the *MYB68* reporter was additionally active in the proximal meristem (**Figure 6A**). Here, PPE cells expressed *MYB68* immediately after the cortex-endodermis initial daughter cells (CEID), while XPE and NPE cells had a significant delay in onset of expression (**Figure 6A, 6B, S3A and S3B**). This radial difference in transcription could be dose-dependently repressed by increasing concentrations of BAP (**Figure 6C and S3C)**. One additional observation was that the onset of *MYB68* expression shifted towards the distal meristematic cells as the root aged (**Figure 6D, 6E, S3E and S3F**). Taken together, we conclude that besides a likely role in controlling the decision to suberize locally in the individual cells, MYB68, or its transcript, likely also influence mechanisms related to periclinal divisions giving rise to MC formation in the meristem.

**Figure 6.**
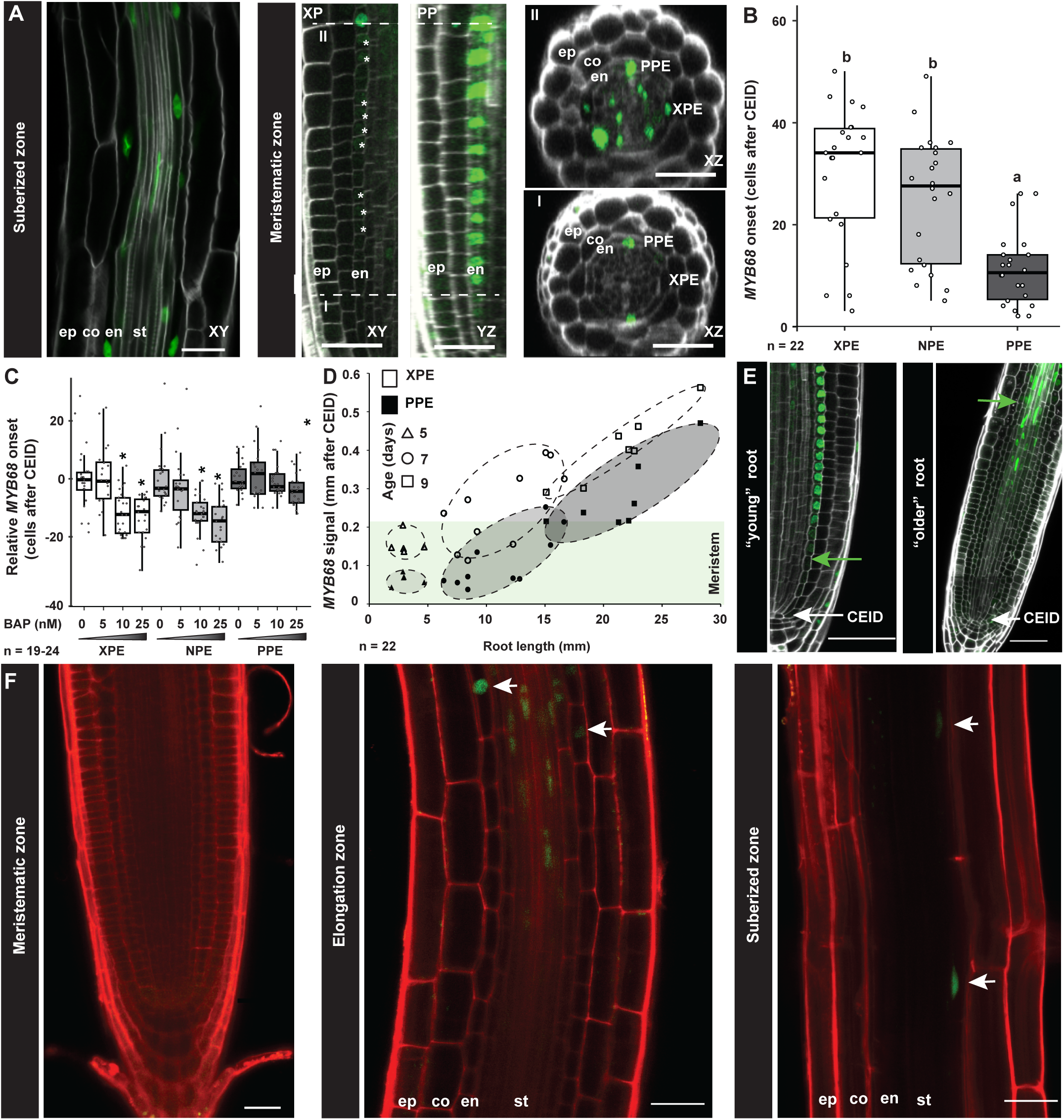
| Analysis of MYB68 localization and expression. A) Activity of MYB68 in roots of 6-day-old seedlings expressing a transcriptional reporter for MYB68 activity (pMYB68:NLS3xVenus). Cell walls were stained using calcofluor white (Ursache, et al., 2018). Scale bar represents 50 µm. The two dashed lines mark the position of cross sections I and II. B) Analysis of expression onset of MYB68 in the meristematic zone of plants from A. Individual letters show signifi-cance according to a Kruskal Wallis test with post hoc Nemenyi test. C) Onset of MYB68 expression in roots of plants grown in the presence of increasing amounts of the artificial cytokinin 6 Benzyl-aminopurine (BAP) normalized to the average expression in mock-treat-ed control roots in each radial position. Asterisks indicate significant differences compared to the untreated sample according to unpaired Wilcoxon text (* p < 0.05). D) MYB68 expression onset in the endodermis. E) Images depicting MYB68 expression start in young (0-6-day-old) and older (7-9 day-old) roots Arrows marks the cortex/endodermal initial daughter cell (CEID). Scale bar represents 50 µm. F) Localization of MYB68 (pMYB68:GFP-MYB68 in myb68-2). Green signal depicts GFP-MYB68 and red propidium iodide (PI). Scale bars represent 25 µm. XPE; Xylem pole associated endodermis, PPE; Phloem pole associated endodermis, NPE; non-pole associated endodermis. Ep; epidermis, co; Cortex, CEID; Cortex endodermis initial daughter cell, en; Endodermis, st; Stele. XP; Xylem pole, PP; Phloem pole.

## Discussion

In this work, we find evidence that the MYB-class transcription factor MYB68 functions in a genetic network connected to radial establishment of endodermal suberization. This is an extension of a recent observation that proposed MYB68 to directly activate expression of suberin-related genes in the periderm (Molina, et al., 2024). However, deeper analysis revealed MYB68 is not only active in endodermal cells that undergo suberization, it additionally accumulates in the nuclei of cells that are outside the area of suberization, i.e. in the elongation zone (**Figure 6**). Thus, in these cells, MYB68 could either indirectly regulate suberization or provide developmental control of endodermal differentiation leading to initiation of the suberization process. This is in line with a recent study that fund evidence for MYB68 working as a higher-tier regulator of other MYB factors that directly influence suberization (Xu, et al., 2022). As MYB factors can participate in protein complexes (Millard, et al., 2019), it is plausible that MYB68 has distinct sets of interaction partners in different tissues, which allow it to provide different functional outputs dependent on the tissue context. Such as system would allow MYB68 to play roles in differentiation, cell division as well as suberization by partaking in distinct regulatory assemblies in its different expression domains.

Intriguingly, our analyses connects the role(s) of MYB68 to a previously established model for PC formation in the proximal meristem. In this model, hormone-based mechanisms that enable and maintain xylem and phloem identities in the vasculature (De Rybel, et al., 2016) extend to the endodermis. This vascular-associated imprint guides PPE cells to a differentional fate that ends in suberization and permit XPE (and NPE) cells to initiate a different trajectory (i.e. PC formation) (Andersen, et al., 2018). Early onset of *MYB68* activity in the PPE cells would thereby serve a non essential role in reinforcing the trajectory that leads to suberization, while delayed expression the early XPE permit patterning events in the early cells that may be necessary for distinct differentiation of these cell files. Since we were unable to detect MYB68 accumulation in the meristematic region, such a regulatory role may not occur at a protein-level, or alternatively, MYB68 could be rapidly degradated in these cells. In favour of the latter, MYB factors related to hormonal signaling have been described as targeted for swift turn-over (Lee and Seo, 2016).

One additional observation was that *myb68* roots has earlier onset of MC formation in the early meristem than Col-0 (**Figure 5**). Thus, similar to what has been proposed in the periderm (Molina, et al., 2024), MYB68 likely also function to suppress periclinal divisions in the young ground tissues. This ties in to the early PPE-biased expression in the meristem, as this provides an inhibitory function in these cell files, which connects with the previous observation that onset of MC formation is specific to the XPE (Baum, et al., 2002). Intriguingly, we further found evidence that onset of *MYB68* activity in the meristem is age-dependent and eventually disappear from the meristematic endodermis (**Figure 6D**). This correlates with the timing of which plants initiate MC formation (Baum, et al., 2002) as well our observed changes in suberin patterning of the endodermis (**Figure 2B**). Intrigingly, the Arabidopsis meristem was recently shown to contain juvenile and adult phases (Yang, et al., 2024). It is intriguing to speculate that this may include a MYB68-containing regulatory network that integrates meristem age, XPE divisions and suberization pattern in the mature root parts, based on meristematic age progression (**Figure 7**).

**Figure 7.**
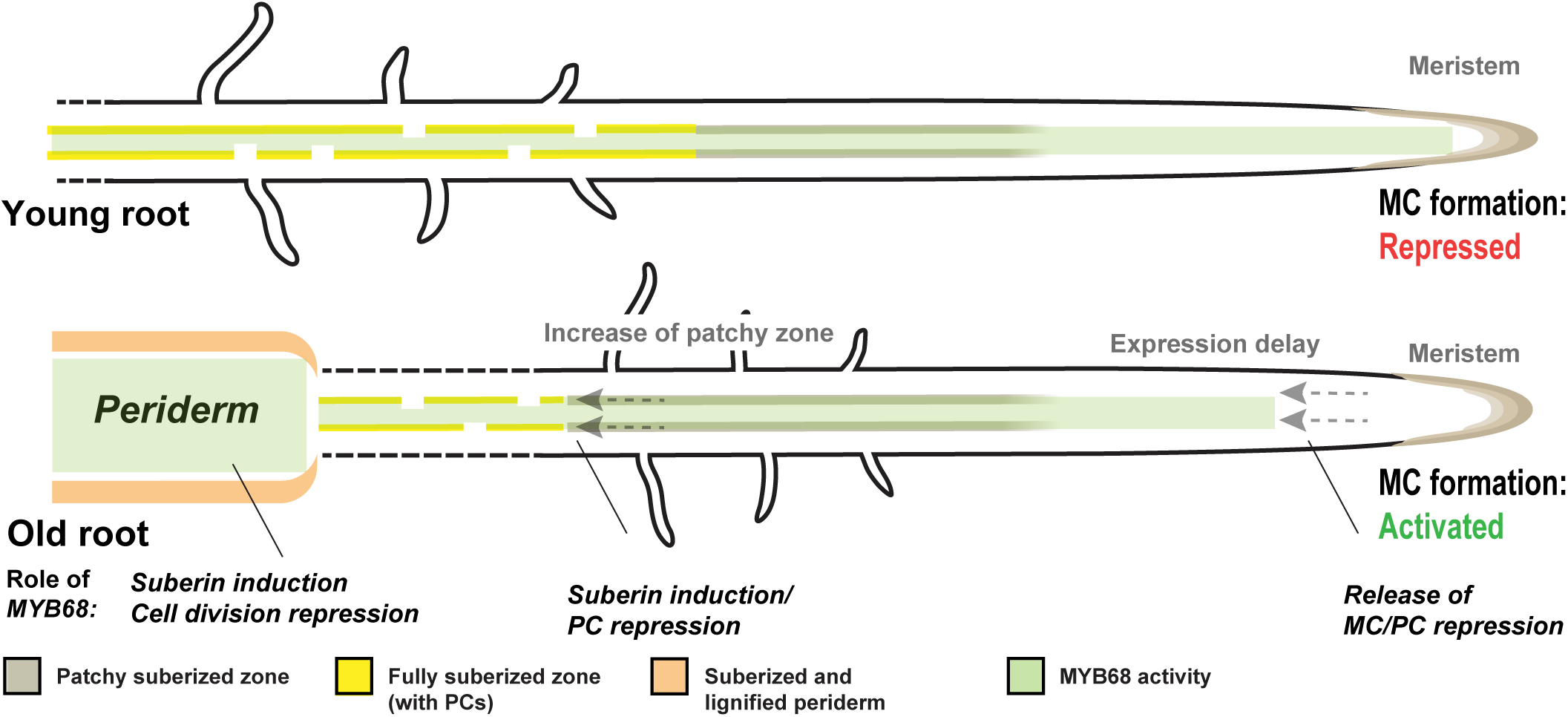
| A model of the role of MYB68 in roots at different developmental stages. In young roots (up to 7-days), MYB68 shows activity in both meristematic and suberizing regions. In the meristem, it represses periclinal cell divisions in the endodermis, which prevents formation of middle cortex cells (MC) as well as passage cells (PC). In the older parts, MYB68 is involved in controlling suberization by regulation of the suberization machinery. As the root ages, absence of the repressing effect of MYB68 on periclinal division in the meristematic region enables MC formation in the xylem pole, this release of repression correlates with a delay in suberiza-tion and suggests a connection between MC and PC formation via MYB68. Moreover, as the root undergoes radial thickening and for periderm, MYB68 serves an additional role in this tissue to control suberization and cell division.

The observation that PC occurrence is increased in *myb68* roots additionally allowed us to probe the function of these elucive cell types. Our finding that several tranporter-coding genes is associated with PC occurrence corroborates a role in nutrient-related transport processes (Holbein, et al., 2021; Lin, et al., 2009; Hamburger, et al., 2002; Gaymard, et al., 1998). Since our identified candidates included transporters related to K+/Na+ homeostasis, PCs (or non-PPE cells) PCs may serve a function in cation homeostasis. As MC formation is increased upon abiotic stress conditions (Cui, 2015), MYB68 may serve and integrative function, which coordinates activation of developmental mechanisms designed to alleviate such conditions for the root. *HKT1*, *KUP8* and *HAK5* were all expressed at the junction between root and hypocotyl, which showed less frequent suberin depositon in Col-0 roots and forms a second cortex layer (Scheres, et al., 1994) (**Figures 5D and S1C**). It is therefore tempting to speculate that this zone may serve to store toxic ions in the ground tissues that are shed during secondary growth and periderm formation. By loading toxic compounds from the vaculature via PCs into cortex cells, this zone could allow the root to expel unwanted ions by directing them into cells destined for expulsion upon periderm formation. While this requires deeper analysis such mechanisms may have important and overlooked implications for root function in cation homeostasis and abiotic stress tolerance.

## Figure legends

**Figure S1.**
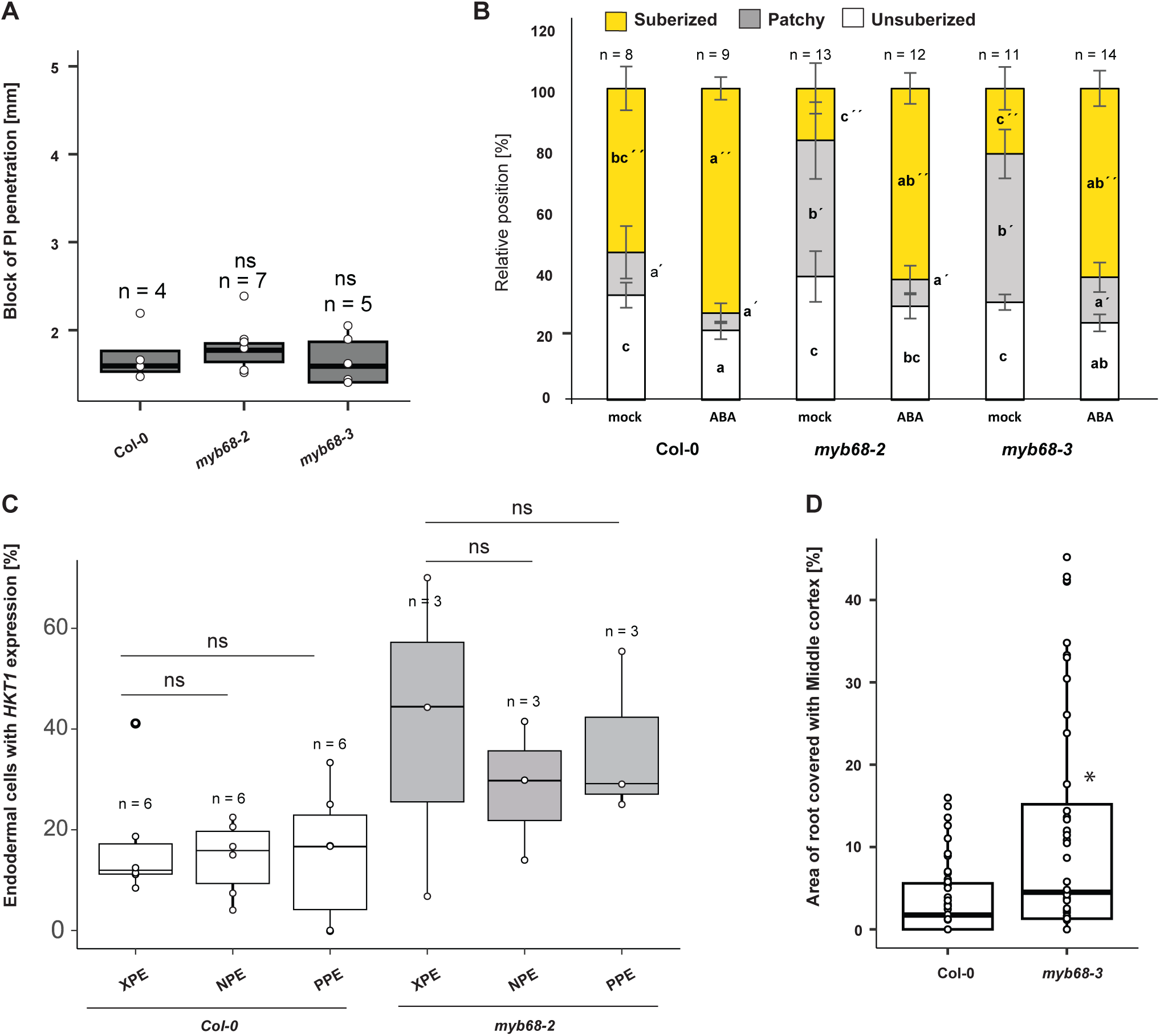
| Analysis of additional *myb68* mutant lines. (A) Functional analysis of CS by measuring onset of propidium iodide (PI) diffusion blockage into the stele (Naseer, et al., 2012). Significant difference vs Col-0 was done according to an unpaired Wilcoxon test. (B) Staining of suberin using the dye Fluorol yellow (FY). Transfer to 1µM ABA or mock 2 days before staining. Individual letters show significance according to the Kruskal Wallis test with the post hoc Nemenyi test. Letter without a prime group unsuberized zones, letter with one prime group patchy zone and letter with two primes group fully suberized zone. (C) Proportion of endodermal cells with expression of *HKT1* in each radial position at ca. 80 % up the longitudinal root axis. Each data point represents the percentage within each analyzed root. A minimum of 5 cells per root per position were quantified. Col-0 and *myb68-2* comparisons in each longitudinal position according to unpaired Wilcoxon test are listed. (D) Measurement of middle cortex (MC) occurrence in Col-0 *vs. myb68-2* in 6-day-old roots grown under mesh conditions. The graph depicts the percentage of root covered with MC measured by measuring the distance from the first-occurring MC to the root tip. Asterisks indicate significant difference to Col-0 according to unpaired Wilcoxon text (* p < 0.05). XPE; Xylem pole associated endodermis, PPE; Phloem pole associated endodermis, NPE; non-pole associated endodermal cells.

**Figure S2.**
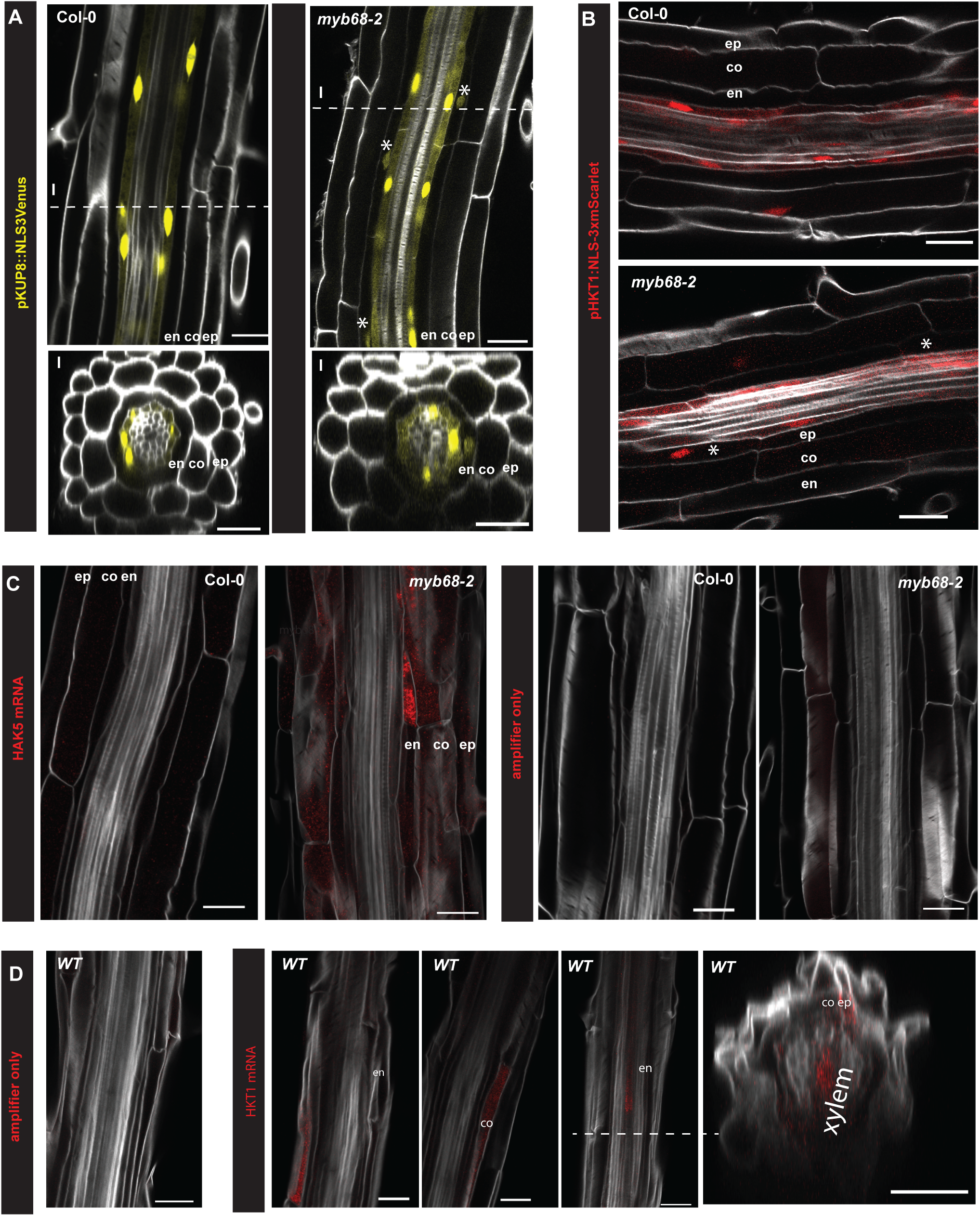
| Localization of transporter associated to passage cell occurrence. A) and B) Expression of *KUP8* and *HKT1* in Col-0 and *myb68-2* backgrounds using the corresponding transcriptional marker lines (*pKUP8*:NLS3xVenus, *pHKT1*:NLS-3xmScarlet). C) For *HAK5* mRNA, fluorescence *in situ* hybridization was employed. As negative control either a probe specific for the Bacillus subtilis dapB with the corresponding amplifier or only the amplifier used for *HAK5* without a probe were used. D) Confirmation of *HKT1* expression outside of the stele in WT (pGPAT5:mCitrine-SYP122). mRNA, fluorescence *in situ* hybridization was employed. As negative control, amplifiers without a probe were used. All images are obtained in the zone of full suberization. Asterisks depict endodermal cells with expression of the respective marker. Cell walls were stained using Calcoflour white (grey signal). Scale bar represents 25 _µ_m.

**Figure S3.**
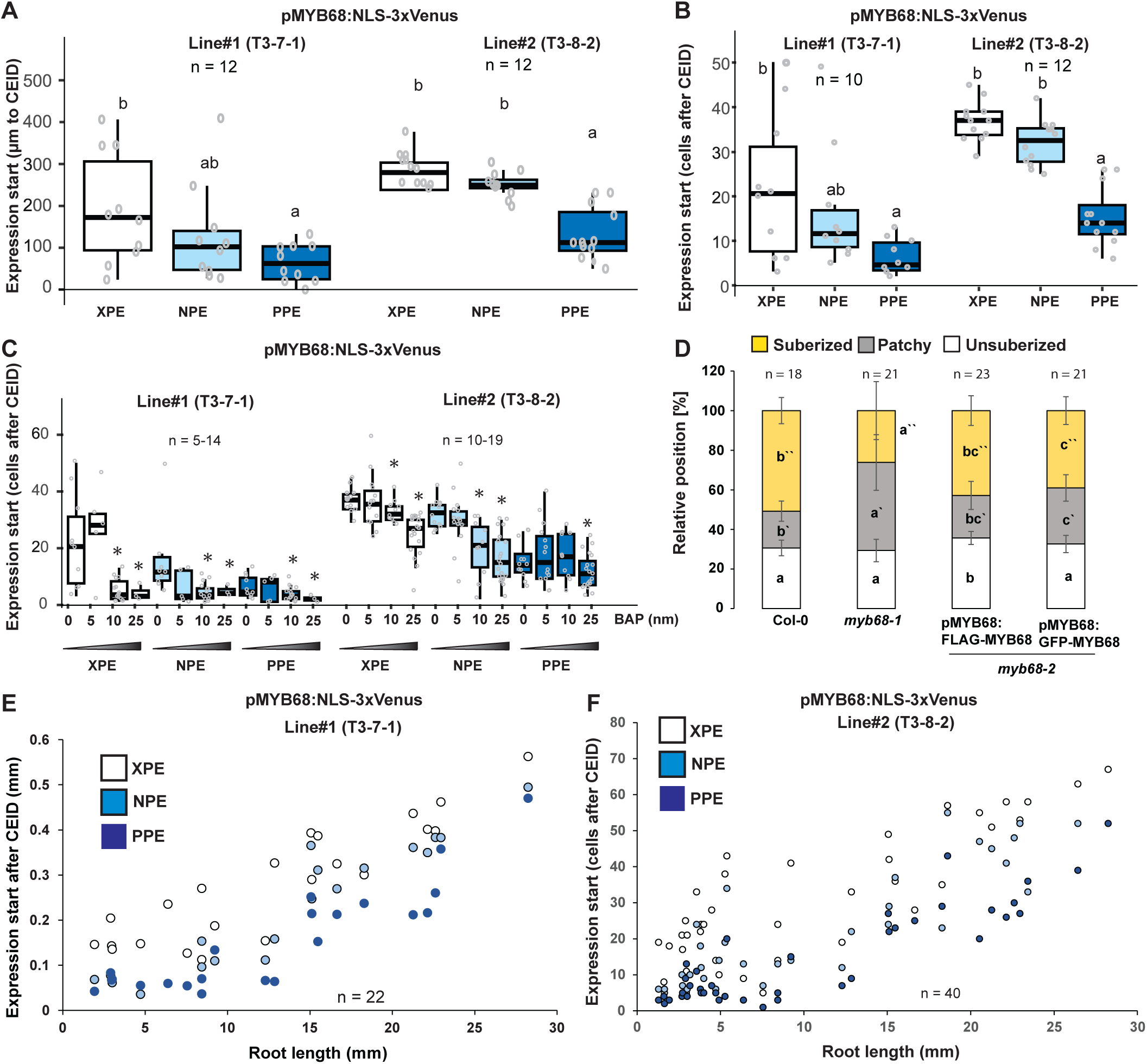
| Expression analysis of *MYB68*. (A) Expression onset of *MYB68* in the meristematic region measured by a transcriptional reporter based on the entire intergenic region upstream of *MYB68* driving expression of a nuclear localized 3x GFP fusion (pMYB68: NLS-3xVenus) (B) Initiation of expression of *MYB68* in each radial position in meristematic endodermal cells the two independent transcriptional marker lines from A. Measurements were taken of the same roots as in A. (C) *MYB68* expression starts along a cytokinin gradient (BAP). Asterisks indicate significant difference to untreated sample according to unpaired Wilcoxon test (P<0.05). (D) Staining of suberin in 6-days-old roots using the dye Fluorol yellow (FY). Percentage of the total root length assigned to zones according to the suberization status in Col-0, *myb68-2*, and two *myb68-2* complementing lines *pMYB68:GFP-MYB68* and *pMYB68:FLAG-MYB68* grown on 1/2MS plates containing a mesh filter. Letters without a prime group unsuberized zones, letters with one prime group patchy zone and letters with two primes group fully suberized zone. (E,F) *MYB68* expression start (E) or number of cells (F) in relation to the cortex endodermal initial daughter cell (CEID). Individual letters show significance according to Kruskal Wallis test with post hoc Nemenyi test or Anova with post hoc Tukeýs HSD test. BAP; 6-Benzylaminopurine, XPE; Xylem pole associated endodermal cells, PPE; Phloem pole associated endodermal cells, NPE; non-pole-associated endodermal cells.

**Table S1: Linear regression analysis of individual suberin zones against root length along a time course**

**Table S2: List of DEGs up in *myb68-2* vs Col-0**

**Table S3: List of DEG up in *myb68-2* vs Col-0 and ns in *myb68-2*vs. Col-0**

**Table S4: Expression of known potassium and sodium transporter genes in *myb68-2* vs. Col-0 and *myb68-2* BAP vs. Col-0**

**Table S5: Expression of known gibberellic acid metabolism or signaling genes in *myb68-2* vs Col-0 and *myb68-2* BAP vs Col-0**

**Table S6: Primers used in this study**

## Materials and methods

### Plant growth

In this study Arabidopsis ecotype Columbia-0 was used. Plants were grown, if not otherwise stated for 5-7 days at long day (16h) and 21°C in 140 _µ_mol/m^2^/s on ½ MS (Duchefa) 0.8 % agar (Bacto Agar) 5.8 pH. For cytokinin treatment, seeds were germinated on ½ MS plates, containing 5,10 or 25 nM N^6^ Benzyladenine (CAS #1214-39-7). For abscisic acid (ABA) treatment 5–day–old seedlings were transferred to ½ MS plates containing 1 _µ_M ABA (CAS # 21293-29-8) and grown for 2 days.

### Cloning

To generate the transcription reporter lines, the corresponding promoters (amplified using primer listed in Table S6) were inserted into a modified pUC19 entry vector (Andersen, et al., 2021) via Infusion cloning (Takara) according to the manufacturer description. The coding sequences of the fluorescent protein and the nuclear localization signal or MYB68 were inserted into a pDONR221 entry vector using BP clonase II (Invitrogen) and recombined using LR clonase II (Invitrogen) into the destination vector pED97 containing a FastRed selection (Andersen, et al., 2018). All constructs were transformed into plants using a modified floral dip method (Logemann, et al., 2006) and selected for FastRed (Shimada, et al., 2010) before propagating further. All mutants in MYB genes were described previously (Molina, et al., 2024).

### Staining procedures

For overall suberin pattern analysis, seedlings were stained in 0,01 % Flourol Yellow 088 solution (w/v) (interchim) in lactic acid for 30 minutes at 70°C, washed and counterstained in a 0,5% Aniline blue solution (CAS # 66687-07-8) (w/v) in water for 30 minutes at room temperature (Naseer, et al., 2012; LUX, et al., 2005). For radial suberin pattern analysis seedlings were fixed in 4% paraformaldehyde (PFA) (CAS # 30525-89-4), cleared with clearsee (10% xylitol (w/v) (Roth CAS 87-99-0), 15% sodium deoxycholate (w/v) (Thermo scientific CAS # 302-95-4) and 25% urea (w/v) (Roth CAS # 57-13-6) and stained with 0.01% Fluorol Yellow dissolved in Ethanol as described previously (Sexauer, et al., 2021). For Casparian strip integrity analysis, seedlings were incubated in a 10 µg/ml propidium iodide (Sigma Aldrich CAS #25535-16-4) dissolved in water and washed with water before imaging (Naseer, et al., 2012). For analysis of transcriptional reporter lines, cell length and middle cortex quantification, seedlings were fixed, cleared and stained with 0.1 % calcofluor white (Fluorescent Brightener 28 CAS #4404-43-7) dissolved in clearesee (Ursache, et al., 2018).

### Transcriptional analysis

For transcriptomic analysis of wild-type and *myb68-2* roots, plants were grown in the presence of mock (DMSO) or 25 nM BAP for 6 days under long day conditions. Total RNA from pooled roots was extracted using a Trizol (Invitrogen) adapted ReliaPrep RNA extraction kit (Promega, Z6012) and subjected to quality control, library preparation, and sequencing on the Illumina NovaSeq platform (Novogene). Approximately 25,000,000 reads (150-bp paired-end), on average, were obtained per sample. Raw reads were processed and cleaned up using fastp (Chen, et al., 2018) with default settings mapped to *A. thaliana* Col-0 genome with the latest annotation (Cheng, et al., 2017) using HISAT2 (Kim, et al., 2019) with default parameters for paired-end reads. Reads per gene without consideration of splicing variants were counted by featureCounts (Liao, et al., 2013) with default parameters. Subsequent statistical analyses were performed using R software (https://www.r-project.org/) unless described otherwise. Differential expression analysis was performed using the *edgeR* package (Robinson, et al., 2009). Library size was normalized by the weighted trimmed mean of M-values (TMM) method, and normalized read counts were fitted to a generalized linear model (GLM) with a negative binomial distribution to identify significantly DEGs. GO enrichment analysis was performed by Metascape (Zhou, et al., 2019).

### Fluorescent in situ hybridization

In situ hybridization chain reaction was performed according to (Oliva, et al., 2022). In short, 6-day old roots were fixed with formaldehyde (CAS # 50-00-0) and dehydrated with a series of increasing concentrations of ethanol and incubation at 100% methanol at −20°C overnight. After rehydration with a decreasing series of methanol concentrations, cell wall was digested in a solution containing cellulase, pectolyase, macerozyme and pectinase at room temperature for 15 minutes. After proteinase K incubation and additional fixation steps, probe hybridization (*HAK5* probe LOT # RTI354, Molecular Instruments) was performed overnight at 37°C. After washing, 10 μl of 3 μM of B3h1 (LOT#S073325, Molecular Instruments) and H2 (LOT#S075725, Molecular Instruments) hairpin solution was added in amplification buffer, allowing amplification at room temperature for 16 h. As negative controls either a probe specific for the Bacillus subtilis dapB gene (LOT # RTB 421, Molecular Instruments) with the corresponding amplifier B1h1 (LOT #S046325, Molecular Instruments) and B1h2 (LOT #S050925, Molecular Instruments) or just the amplifier used for *HAK5* B3h1 (LOT#S073325, Molecular Instruments) and B3h2 (LOT#S075725, Molecular Instruments) without a probe were used. Excess hairpin was washed away, seedlings were cleared with clearsee (10% xylitol (w/v) (Roth CAS 87-99-0), 15% sodium deoxycholate (w/v) (Thermo scientific CAS 302-95-4) and 25% urea (w/v) (Roth CAS # 57-13-6) and stained with calcofluor white (Fluprescent Brightener 28 CAS #4404-43-7).

### Microscopy/imaging

Endodermal suberin patterning was imaged with an AxioZoom V16 using a standard GFP filtercube. The parts of the root was defined as follows: Unsuberized zone: from root tip to first suberized cell; patchy zone: from first suberized cell to the first point where all cells across the circumference of the endodermis are suberized; fully suberized zone: from first ring of suberized cells to the root-hypocotyl junction. Suberin radial pattern, middle cortex, transcriptional marker lines and FISH samples were imaged with a confocal laser scanning microscope (Zeiss LSM 980) with the following settings: flourol yellow (ex: 488 nm, em: 500-550 nm), propidium iodide (ex: 561 nm, em: 600-650 nm), calcofluor white (ex: 405 nm, em: 407-466 nm),Venus transcriptional marker line (ex: 514 nm, em: 500-560m), Scarlett signal (ex: 561 nm, em: 570-640 nm), FISH samples (ex: 639 nm, em: 650-680nm). Confocal images were taken with the 40x (NA 1.2) oil immersion objective and Zen Connect was used to determine the exact position of the image in the root.

### Statistics

Statistical analysis was performed in R. Data sets were analyzed for normal distribution using the Shapiro-Wilk test. In case data was normally distributed One-way Anova with post hoc Tukeýs HSD test was performed for multiple comparisons and Students T test or Welch two sample test was performed for comparison of two groups. In case data was not normally distributed for multiple comparison Kruskal Wallis test with post hoc Nemenyi test was used and for comparison of two groups unpaired Wilcoxon test was employed. For time-course experiments linear regression analysis was conducted. All analysis were performed at least twice in independent experiments.

### Radial suberization pattern analysis

We constructed one-dimensional arrays for each cell file, and assigned the values 1 and 0 to each suberized and unsuberized cell, respectively. We first normalized the length of all arrays to an arbitrary size (we choose *N = 2000* units), a procedure that allows the direct comparison between cell files and roots with different cell number. After this rescaling, all arrays have the same length, but the size of the corresponding cells has changed (Figure 2A). To analyse the progression of suberization along different cell files, we computed the normalized cumulative sum *CS_i_* of the rescaled arrays, defined as 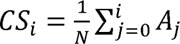, where *CS_i_* is the *i*-th element of the cumulative sum, *N* is the length of the array, *A_j_* are the entries of the suberin arrays (e.g. *A_j_* = [0,0,1,0,1, . . ., 1,1,1]), and index *j* runs from 0, the start of the array, to the *i*-th element of the array. The cumulative sum progressively adds the contributions of every suberizedcell, reaching a final value that will depend on the number of 1s and 0s in the array. For instance, for a completely suberized root, the normalized cumulative sum will take the form 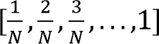, reaching a maximum value of 1. The cumulative sum is a strictly non-decreasing function, and its final value represents the proportion of suberized cells of an array.

## Acknowledgments

All authors thank Ton Timmers and the Central Microscopy facility (CeMic) for microscopy aid. Aristeidis Stamatakis and his greenhouse team at MPIPZ are thanked for help with plant growth. BDR and APM are thanked for tea and coffee, respectively.

## Author contributions

Conceptualization: TGA, LK and LR. Methodology: LK, RTN, DM, LR. PFJ, JMM and TGA. Investigation:LK, RTN and TGA. Visualization: LK, PFJ, JMM and TGA. Funding acquisition: TGA, LR. Main writing: TGA and LK. All authors read and commented on the final version of the manuscript.

## Data and materials availability

RNA-seq raw reads generated in this study have been deposited at National Center for Biotechnology Information under BioProjectID: PRJNA1110212

